# imageseg: an R package for deep learning-based image segmentation

**DOI:** 10.1101/2021.12.16.469125

**Authors:** Jürgen Niedballa, Jan Axtner, Timm Fabian Döbert, Andrew Tilker, An Nguyen, Seth T. Wong, Christian Fiderer, Marco Heurich, Andreas Wilting

## Abstract

1. Convolutional neural networks (CNNs) and deep learning are powerful and robust tools for ecological applications. CNNs can perform very well in various tasks, especially for visual tasks and image data. Image segmentation (the classification of all pixels in images) is one such task and can for example be used to assess forest vertical and horizontal structure. While such methods have been suggested, widespread adoption in ecological research has been slow, likely due to technical difficulties in implementation of CNNs and lack of toolboxes for ecologists.
2. Here, we present R package imageseg which implements a workflow for general-purpose image segmentation using CNNs and the U-Net architecture in R. The workflow covers data (pre)processing, model training, and predictions. We illustrate the utility of the package with two models for forest structural metrics: tree canopy density and understory vegetation density. We trained the models using large and diverse training data sets from a variety of forest types and biomes, consisting of 3288 canopy images (both canopy cover and hemispherical canopy closure photographs) and 1468 understory vegetation images.
3. Overall classification accuracy of the models was high with a Dice score of 0.91 for the canopy model and 0.89 for the understory vegetation model (assessed with 821 and 367 images, respectively), indicating robustness to variation in input images and good generalization strength across forest types and biomes.
4. The package and its workflow allow simple yet powerful assessments of forest structural metrics using pre-trained models. Furthermore, the package facilitates custom image segmentation with multiple classes and based on color or grayscale images, e.g. in cell biology or for medical images. Our package is free, open source, and available from CRAN. It will enable easier and faster implementation of deep learning-based image segmentation within R for ecological applications and beyond.

## Introduction

Technological advances over the last decades have led to unprecedented increases in the amount of data available for ecological research, which require efficient and reproducible analyses (Christin, Hervet, & Lecomte, 2019). Methodological developments in machine learning paired with improvements in computational power enabled an increasingly widespread use of deep learning methods to make use of these data in a multitude of ecological applications, particularly for visual recognition (Kattenborn, Leitloff, Schiefer, & Hinz, 2021).

Convolutional neural networks (CNN) have been at the core of many of these innovations thanks to their ability to analyze complex nonlinear data and performance at near-human or even exceeding human levels in specific tasks (Grace, Salvatier, Dafoe, Zhang, & Evans, 2018; Mnih et al., 2015). CNNs are multi-layered artificial neural networks that consist of an input layer, stacked hidden layers (for example convolutional and pooling layers), and the final output layer (Minaee et al., 2021). The number of the hidden layers defines the depth of the network and this is where filters introduce translation invariance and parameter sharing between pixels. These filters set every pixel in relation to its surrounding pixels, producing so-called feature maps which are then pooled (LeCun, Bengio, & Hinton, 2015).

One prominent domain of CNNs is image segmentation, which classifies each pixel of an image, providing locality information for each label class. Here, we primarily focus on two ecological applications of image segmentation in the context of forest structural metrics from color photographs, the assessment of i) tree canopy density (canopy closure and canopy cover), and ii) understory vegetation density. Moreover, we provide an outlook on how the package imageseg can be used for grayscale images or more complex multi-class applications.

Canopy density and understory vegetation density are important metrics of forest structure and are therefore important for a number of ecological applications from biodiversity surveys to forest monitoring and ground-truthing remote sensing data. Canopy density is related to canopy architecture, forest conditions, carbon and stand densities, and affects light regimes and microhabitat structure (Jennings, Brown, & Sheil, 1999). Understory vegetation is composed of immature trees, shrubs, and herbaceous vegetation, and is thus linked to forest regrowth and plant succession (e.g. after disturbances), while also providing essential resources and shelter for terrestrial wildlife (McShea & Rappole, 1992; Nilsson & Wardle, 2005). Both measures are thus related to carbon sequestration and storage, biodiversity conservation, and ecosystem regeneration. Furthermore, due to their importance for forestry and crop growth, both measures are also economically relevant.

Traditionally, tree canopy density was often assessed using manual methods such as spherical densiometers or photometric techniques requiring specialist equipment (see Paletto & Tosi, 2009 for comparison of methods). Traditional methods for quantification of understory vegetation density included vegetation density boards or profile boards (Nudds, 1977). These methods are slow, labor-intensive, and often require special equipment. Moreover, the manual approaches are often non-reproducible, can be subject to observer bias and dependent on experience. Remote sensing technologies such as airborne or terrestrial laser scanning have been successfully used to determine forest structural attributes (Campbell, Dennison, Hudak, Parham, & Butler, 2018; Latifi et al., 2016; S. Li et al., 2021; Wing et al., 2012; Zong, Wang, Skidmore, & Heurich, 2021). These methods, however, whilst powerful and accurate, remain constrained by logistical challenges and often prohibitive costs (Hummel, Hudak, Uebler, Falkowski, & Megown, 2011).

Deep learning methods have been developed to help address these shortcomings and streamline analyses of forest structural metrics using easy-to-collect digital photographs (Abrams et al., 2019; Díaz, Negri, & Lencinas, 2021; K. Li et al., 2020). The approaches outlined in Abrams *et al*. (2019) and Li *et al.* (2020) apply image segmentation based on the U-Net architecture introduced by Ronneberger *et al.* (2015), whereas the model of Diaz *et al*. (2021) uses deep learning regression to assess plant area index (PAI) and is thus not an image segmentation approach (i.e. it does not retain pixel-level information). These deep learning methods achieve higher accuracy than manual methods whilst being much faster and more cost-efficient (Abrams et al., 2019; K. Li et al., 2020). Uptake of deep learning methods in ecological applications, however, was constrained by technical difficulties in implementation and the lack of a readily available workflow or software package to easily apply these methods. Furthermore, training data sets were often relatively homogeneous (i.e. from a single forest ecosystem), leaving doubts about how applicable such models are across ecosystems.

Here, we present the R package imageseg, which implements a complete workflow for binary and multi-class image segmentation using deep learning models implemented in TensorFlow (Abadi et al., 2015). It works with grayscale as well as color input images. We illustrate the workflow by providing two new models for canopy density and understory vegetation density. Both models were built using the largest and most diverse data set for these applications to date. The canopy density model is the first to be explicitly trained on both canopy closure and canopy cover images. Both models yielded highly accurate predictions. Beyond these forestry-related applications, the package and its workflow can be adopted for the development of custom image segmentation tasks in other fields, e.g. in microscopy or cell biology.

The imageseg R package presents a critical advancement in deep learning-based image segmentation. While imageseg is particularly relevant to applied research, its accessibility promises to facilitate image segmentation across disciplines. The pre-trained models for canopy and understory vegetation density, along with the simple workflow of imageseg, will enable faster and more reproducible assessment of important forest structural metrics from easily obtainable data. We believe that imageseg will have numerous applications relevant to forestry, ecology and conservation, but will also offer solutions across scientific disciplines.

## Package description

### Overview

We present imageseg, an R package which implements a general-purpose image segmentation workflow based on convolutional neural networks using the U-Net architecture (Ronneberger et al., 2015). It is suitable for binary and multi-class image segmentation problems based on grayscale and color images. Models are implemented in R via Keras (Chollet & et al., 2015; Kalinowski et al., 2021) using a TensorFlow backend (Abadi et al., 2015).

The workflow covers data processing, model training, and predictions for image segmentation tasks applied to digital images. In total, the package contains eight main functions for image processing, model creation, predictions, as well as the extraction of relevant information from predictions (see Table 1). The package workflow is explained in a detailed vignette and illustrated in Figure 1.

**Table 1:** List of functions of the imageseg package.

| Function | Description |
| --- | --- |
| *resizeImages* | Resize and save images |
| *findValidRegion* | Subset image to valid (informative) region |
| *loadImages* | Load image files with magick |
| *dataAugmentation* | Image data augmentation: rotating and mirroring images, adjusting colors |
| *imagesToKerasInput* | Convert magick images to keras input (4D arrays) |
| *u_net* | Create a U-Net architecture |
| *loadModel* | Load TensorFlow model and custom objects from hdf5 file |
| *imageSegmentation* | Model predictions from images based on TensorFlow model |

**Figure 1:**
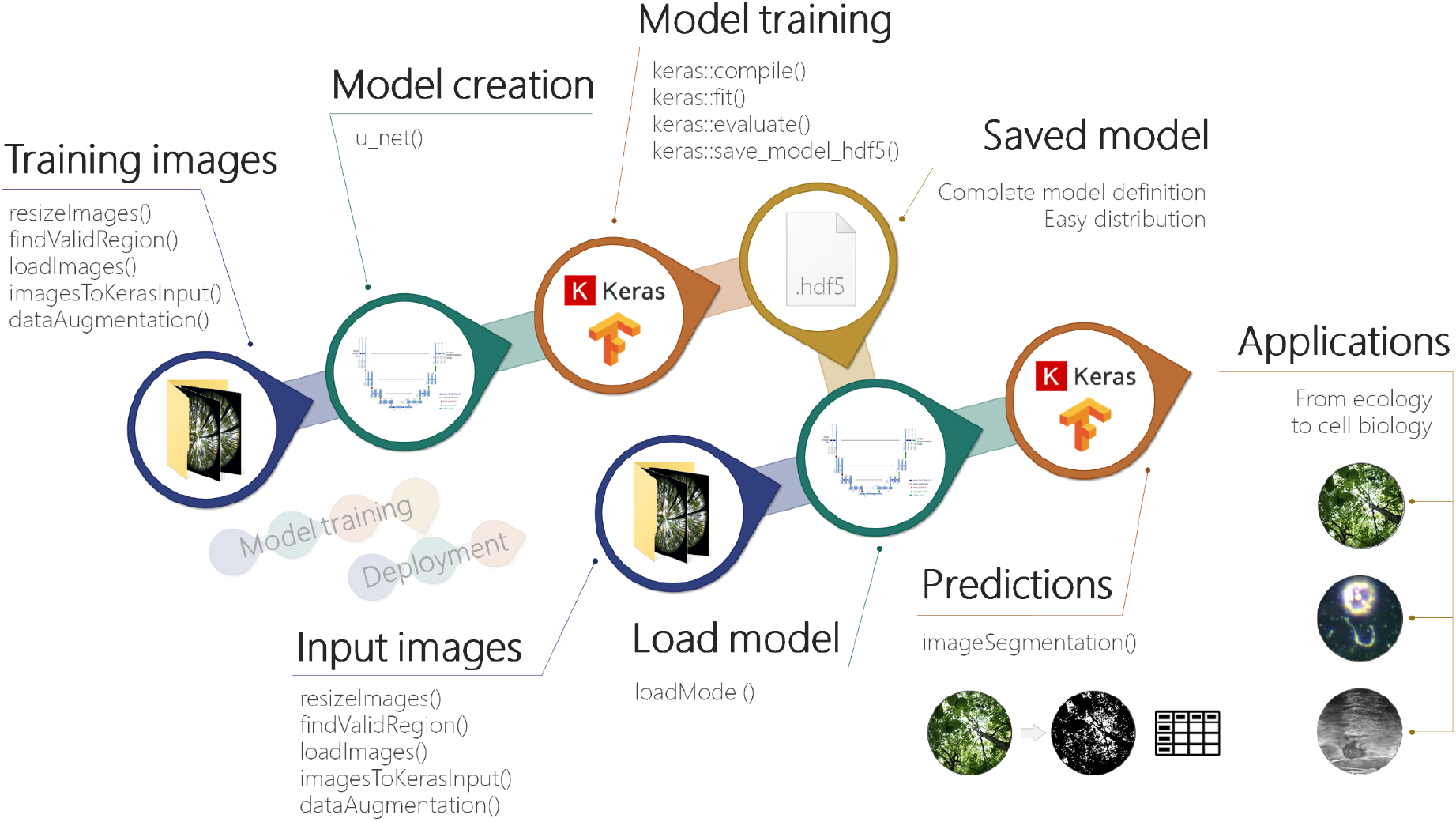
Workflow of the R package imageseg showing the main steps and their relevant functions.The steps at the top show the model training workflow and the steps at the bottom show the model deployment workflow.

Our package is intended for two main purposes:

1. General-purpose model training for image segmentation, including both de-novo training with custom model architectures and continued training of existing models. The package is suitable for both binary and multi-class image segmentation.
2. Model predictions using pre-trained models, allowing users to easily perform image segmentation on digital images. We provide pre-trained models for two types of forest structural characteristics (tree canopy density and understory vegetation density) along with respective training data for download.

Here, we demonstrate the utility of the package for image segmentation and illustrate its application for forest structural metrics, namely a model for canopy density (differentiating canopy and sky), and a model for understory vegetation density. Both of these applications perform binary classifications of all pixels in input images. In addition, the imageseg package also supports multi-class image segmentation (see the example for the detection of bacteria, erythrocytes and background in darkfield microscopy images in Figure 2).

**Figure 2:**
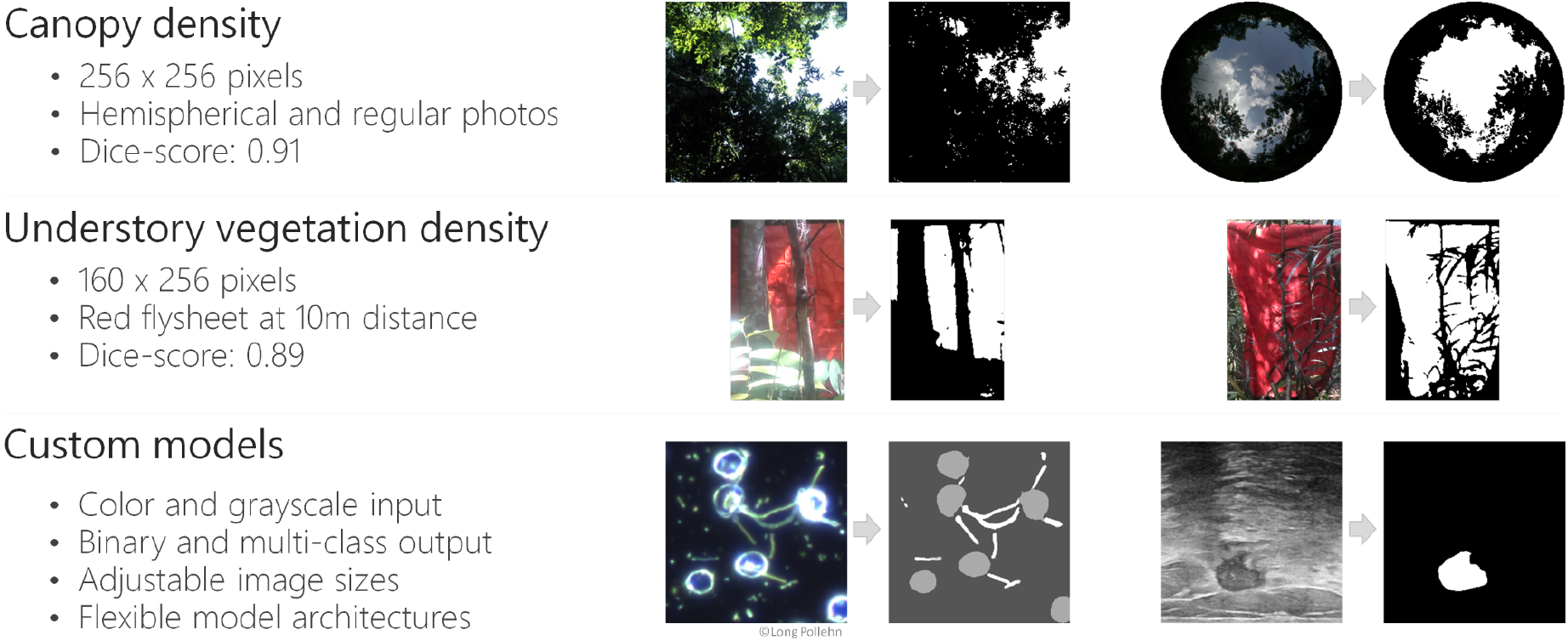
Example output of imageseg. The canopy and understory vegetation density models are provided for download. The package workflow is furthermore suitable for custom image segmentation tasks. The examples at the bottom show a multi-class image segmentation of a (color) microscopy image and a binary segmentation of a (grayscale) ultrasound breast cancer scan image (see package vignette for code and data sources).

### Data processing

The imageseg package accepts jpg, tif and png images as input. Image processing in R is mainly performed by the magick package (the R interface to ImageMagick, Ooms (2021)), allowing for easy viewing of images in R.

Data processing is divided into a logical sequence of steps, implemented in a few functions (see Table 1 and Figure 1). *resizeImages* resizes original images to the dimensions expected by the models while allowing flexibility to prepare input for custom models also. The function can optionally crop images or remove uninformative image regions. The *findValidRegion* function can help identify the valid (informative) image region for cropping of images (e.g. for removing the black, uninformative borders from hemispherical canopy photos taken with fisheye adapters). *loadImages* loads images into R as magick objects, allowing for easy viewing of images in R. *dataAugmentation* performs optional and adjustable data augmentation (rotation, mirroring, modulating brightness, saturation and hue) to increase the amount and variability of available training data and thus the robustness and invariance of the CNN (only relevant for model training, not for model predictions). *imagesToKerasInput* converts the images to model input required by the CNN (4D-array with dimensions: image - row - column - channel).

### Model training

Users who wish to train their own model can create a U-Net based architecture with the *u_net* function, which was slightly modified from the original function in the platypus package (Maj, 2020). It supports flexible image sizes, data types, number of U-Net blocks, layers per block, convolution filters, the optional inclusion of dropout layers, and supports binary and multi-class classifications (thus also allowing general-purpose image segmentation as shown in Figure 2).

As mentioned above, the *dataAugmentation* function can perform various data augmentation tasks prior to model training to increase the amount and diversity of available training data. Users who wish to only apply pre-trained models can skip the model training steps.

### Model predictions

*loadModel* loads a pre-trained model from a hdf5 file (containing the entire architecture and training state of the model in a single file) and ensures the relevant custom objects are loaded (custom loss functions and metrics). *imageSegmentation* is the main function for predictions and performs image segmentation using the model and images provided by the user. It returns the pixel-wise probabilities for the predictions, the classified image segmentation masks based on the input images, and summary tables. In binary predictions (such as the vegetation structure models we present) these summaries represent the desired vegetation metrics (openness / gap fraction) along with their opposite (canopy closure / cover, or understory vegetation density). It allows for adjustable masking of hemispherical canopy images (e.g. circular masks to exclude uninformative corners in hemispherical images). In models with multiple output classes, it returns classified image segmentation masks, probability layers for each class, and summarizes prevalence and probabilities of all predicted classes.

## Vegetation structure models

We demonstrate the utility of the imageseg package for image segmentation of two types of forest structural information: a model for tree canopy density (differentiating canopy and sky in canopy cover images and hemispherical canopy closure images), and a model for understory vegetation density. Both models use color photographs as input and were trained on the largest and most diverse training data set for these forest structural metrics to date.

Here, we follow Jennings, Brown and Sheil (1999) in defining canopy closure as the proportion of sky hemisphere obscured by vegetation when viewed from a single point. It is commonly recorded using canopy hemispherical photography (CHP), e.g. via fisheye lenses. In contrast, canopy cover is the percent forest floor occupied by the vertical projection of tree crowns. It is recorded using canopy cover photography (CCP) (Chianucci, 2016; Macfarlane, Grigg, & Evangelista, 2007). While cameras cannot technically record the exact vertical projection of tree crowns (due to the camera sensor being essentially a single point), the narrower field of view and higher focal length ensure the canopy cover images approximate actual canopy cover well.

We define understory vegetation density analogously to canopy cover as the percent area covered by the lateral obstruction from understory vegetation. It is assessed using a red flysheet of 1×1.5m held at a distance of 10 meters from the camera. The red background color was chosen to provide a strong color contrast with vegetation. The method is similar to the cover board photography method in Campbell *et al.* (2018).

### Model architecture and training

Both models (canopy and understory vegetation density) share the same underlying U-Net architecture (Ronneberger et al., 2015). They use color images as input and only differ in the dimensions of the input images. Both models perform binary classification with one output class. The model architecture is a standard U-Net with 32 feature maps in the first convolution (the number of feature channels doubles at every downsampling step along the contracting path).

During model training we used BCE-Dice loss (sum of binary cross-entropy and soft-Dice loss) and the ‘Adam’ model optimization algorithm (Kingma & Ba, 2017) with the following parameters: initial learning rate = 0.001, Beta_1 = 0.9, Beta_2 = 0.999, decay rate = 0. We reduced the learning rate by a factor of 5 when learning stagnated (with patience 3) and employed an early stopping strategy with a patience of 5 to prevent overfitting, keeping the model weights from the epoch with the lowest validation loss (Caruana, Lawrence, & Giles, 2000). We used a batch size of 12 for the canopy model and 16 for the understory model. Maximum number of epochs was set to 100, but early stopping led to shorter model training.

In all models, 80% of the original images were used for training and 20% of images were used as test data for model evaluation. Training images were subject to data augmentation to increase robustness of the network and invariance (e.g. to lighting conditions and image geometry). Of the augmented training data, 15% were used for model validation. Model accuracy was evaluated using two similarity scores, Dice similarity coefficient and Jaccard index. See Table 2 for details on model training and results.

**Table 2:** Overview of training data and results of the canopy density and understory vegetation density models.

|  | **Canopy model** | **Understory vegetation model** |
| --- | --- | --- |
| Original images | 4109<br>(CHP: 1310, CCP: 2799) | 1835 |
| Training images<sup>1</sup> | 16769 | 2496 |
| Validation images<sup>1</sup> | 2959 | 440 |
| Test images<sup>2</sup> | 821 | 367 |
| Image size (WxH) | 256x256 | 160x256 |
| Data augmentation |  |  |
| rotation | 90°, 180°, 270° | - |
| mirroring | horizontal and vertical axes | vertical axis |
| Dice score | 0.91<br>(CHP: 0.91, CCP: 0.92) | 0.89 |
| Jaccard index | 0.87<br>(CHP: 0.86, CCP: 0.87) | 0.84 |

<sup>1</sup> after data augmentation
<sup>2</sup> without data augmentation (only original images)

Training was conducted in R 4.1.1 with keras 2.4.3 and TensorFlow 2.2.0 on a Windows workstation with 128Gb of RAM and an Nvidia GeForce GTX1660 Super GPU.

### Training data

Compared to previous approaches, our models use a larger and more diverse training data set. Both canopy and understory data have broader geographical scope by including training data from Viet Nam, Laos, Sabah (Malaysian Borneo), and in the case of CCP data, Germany. The canopy model was trained on CCP and CHP images. Images were taken under varying illumination conditions. Image masks were created manually in the GNU Image Manipulation Program (GIMP) (The GIMP Development Team, 2019). See Table 2 and Supporting Information S1 for more details on the training data used in both models and their processing.

#### Canopy model

For the canopy model, we used a total of 4109 canopy photographs (and their respective segmentation masks). Of these, 1310 were hemispherical CHP images (from Sabah, Malaysian Borneo) and 2799 were CCP images (from Malaysia, Viet Nam, Laos and Germany). 69 of the CCP images were sky photos for which we assessed classification accuracy separately.

CHP images cover multiple habitat types along a gradient from primary tropical rainforest, over secondary forest subjected to varying degrees of logging, to oil palm plantations within the Stability of Altered Forest Ecosystems (SAFE) project area in Sabah, Malaysian Borneo (Ewers et al., 2011). CCP images cover primary and secondary tropical rainforests in three countries (along an elevational gradient from 60 m to 1,420 m a.s.l.), and also include temperate coniferous and mixed mountain forests of the Bavarian Forest National Park, Germany (covering an elevational gradient from 600 to 1450 m a.s.l; Cailleret, Heurich, & Bugmann, 2014).

All canopy images were resized to 256×256 pixel resolution while preserving the original aspect ratio (thus slightly cropping the rectangular original images to obtain square images). We increased the number of images available for training and validation via data augmentation, which included image rotation (90, 180 and 270 degrees) and mirroring along the horizontal and vertical axes, resulting in a total of 16769 images for training and 2959 for validation (see Table 2). We used 821 unaugmented images for model evaluation.

#### Understory model

For the understory vegetation density model, we used 1835 images and their respective segmentation masks (866 from Malaysia, 842 from Viet Nam and 127 from Laos). Images were taken of a 1×1.5m red flysheet held at a distance of 10 meters from the cameras and cropped to contain only the extent of the flysheet (and the vegetation covering it). Cropping could not be automated due to high variation in images and was thus done manually (the need for cropping can be avoided if images are taken zoomed in and the flysheet fills the image frame). All understory images were resized to 160×256 pixels (aspect ratio 5:8). Data augmentation included mirroring images along the vertical axis, resulting in a total of 2496 images for training and 440 images for validation. Model evaluation was done with 367 unaugmented images (see Table 2).

### Model results

The canopy model achieved an overall Dice similarity coefficient of 0.91 and Jaccard index of 0.87. The respective scores were 0.91 and 0.86 for CHP images, and 0.92 and 0.87 for CCP images. Both indices were 1 for sky photos. The understory vegetation density model achieved an overall Dice coefficient of 0.89 and Jaccard index of 0.84 (see Table 2).

### Performance

Model predictions can be created quickly with standard hardware and on CPU alone. A laptop with an Intel i7-7500U CPU created predictions for about 100 canopy images per minute. A graphics card is not required, but if available can speed up predictions and model training considerably.

## Application examples

Our pre-trained forest structure models are suitable for quickly deriving habitat metrics from easy-to-collect data using powerful CNNs. The diversity of the model training data sets allow for greater flexibility and generalization strength than previous models. They can be applied to record canopy cover and vegetation density in biodiversity studies (Asad et al., 2020; Tilker et al., 2020), for ground-truthing in remote sensing applications (Hansen et al., 2019; Campbell et al. 2018), or in monitoring of forest pathology, forest conditions or stresses (e.g. under drought conditions). Researchers in crop sciences might also find such models helpful for assessments of crop growth using photographic data.

Furthermore, the package and workflow are not restricted to forest structural metrics and binary classifications, but are universal and can also be applied to image segmentation tasks in other domains, including for multi-class image segmentation (e.g in microscopy or cell biology, see Figure 2). The package and its workflow were designed to be accessible and simple to use in order to facilitate simple and quick experimentation without the need for in-depth technical knowledge about CNNs.

The generalization strength of CNNs ensures that the methods are robust and not dependent on optimal lighting conditions, thus allowing greater flexibility in data collection, e.g. under field conditions where waiting for optimal conditions is often not an option, or for cellular and microscopic images which are often highly variable.

## Conclusion

This paper presents an R package for image segmentation using deep convolutional neural networks based on the powerful and flexible U-Net architecture (Ronneberger et al., 2015). The package implements a complete workflow including image processing, model creation and prediction. It was designed to be accessible and provide a simple and intuitive user experience that does not require specialist-level technical understanding of deep learning models and CNNs. imageseg constitutes a major advancement towards making deep learning-based image classification available to non-specialists in a convenient and easy-to-use R package.

We demonstrate the usefulness of the package for deriving forest structural metrics (canopy and understory vegetation density) from color photographs and provide pre-trained models for these metrics. The models were trained on large and diverse training data sets and achieved high classification accuracy (Dice similarity coefficient of 0.91 for canopy images and 0.89 for understory images). Beyond this immediate application, the workflow and functions we provide are flexible and allow for general-purpose image segmentation, making the package a versatile tool for simple implementation of complex image segmentation workflows in R.

## Data accessibility

imageseg is a free and open source R package. The latest release version is available on CRAN (https://CRAN.R-project.org/package=imageseg) and can be installed via `install.packages(“imageseg”)`. A tutorial vignette is included in the package. Source code and the development version are available from GitHub (https://github.com/EcoDynIZW/imageseg). Links to the pre-trained models, classification examples and data used for model training and testing are available from: https://github.com/EcoDynIZW/imageseg. R code to create the described models is included in Supporting Information S2.

## Acknowledgements

We thank the Sabah Biodiversity Center for issuing research permits and the Sabah Forestry Department for its support. We thank Azlan bin Mohamed, John Mathai, local research assistants and others involved in fieldwork for data collection and processing of canopy cover and understory photos from Sabah. In Vietnam and Laos we thank the staff of the WWF-CarBi project for providing extensive logistical support; Bach Ma NP, the Thua Thien Hue and Quang Nam Saola NRs, and Xe Sap NPA for providing permissions and personnel to conduct this research; and our field team leaders. This project received financial support from the German Federal Ministry of Education and Research (BMBF FKZ: 01LN1301A), the United States Agency for International Development (USAID) through Cooperative Agreement No. 72044020CA00001 “USAID Biodiversity Conservation” Activity, Point Defiance Zoo and Aquarium, San Francisco Zoo and Leibniz-IZW. T.F.D. was supported by a University of Western Australia PhD scholarship and a CSIRO top-up scholarship. We thank the SAFE project, South East Asia Rainforest Research Programme (SEARRP) and IDEA Wild for logistical support and equipment. Marion Pfeifer assisted with canopy hemispherical photo processing. Data collection strongly benefitted from local traditional knowledge. We thank the German Federal Agency for Nature Conservation for funding research in Bavarian Forest National Park (FKZ: 3518830200). The Bavarian Forest National Park thanks all field assistants who took the canopy pictures and processed them. We are grateful to Oliver R. Wearn and Adam F. Smith for their networking efforts which connected the collaboration partners in this project.

## Author’s contribution

J.N., J.A. and A.W. conceptualized the idea. J.N. and J.A. developed the package and conducted model training. T.F.D. provided the canopy hemispherical photography data. A.T., A.N. and S.T.W. provided the canopy cover and understory vegetation training data from Sabah, Vietnam and Laos. C.F. and M.H. provided canopy cover training data from Bavarian Forest National Park. All authors contributed to writing the paper.

