## Supporting Information S1 for "imageseg: an R package for deep learning-based image segmentation"

### Description of the training data sets

Here we provide a description of the data sets used for model training and evaluation. The canopy data sets were recorded as part of independent projects, thus methodologies differ.

#### Canopy hemispherical photography (CHP)

Canopy hemispherical photos (canopy closure images) were collected in the SAFE project area including the Maliau Basin Conservation Area in Sabah, Malaysian Borneo, spanning gradients in land use and habitat degradation. Specifically, they stem from seven areas with different logging intensities, one old-growth forest and one oil palm plantation. Images were taken at > 1500 2x2m study plots along transects to both sides of future forest fragment edges (or hypothetical forest fragment edges in the case of old-growth forest and oil palm plantation sites, see Ewers et al. 2011 for experimental design details). At each plot four images with three different exposure settings (exposure bracketing: -1.3/-1/-0.7EV) were taken. Images were taken with a Nikon Coolpix 4500 camera using a Soligor Fisheye lens. For model training and testing, we randomly selected one out of the three exposure bracketing images.
Image masks were created manually using CAN-EYE software. In total we used 1310 of the canopy closure images (and their corresponding masks) deemed suitable for model training and evaluation.

#### Canopy cover photography (CCP)

Canopy cover photos were taken at vegetation plots in three forest reserves in Sabah, Malaysian Borneo (Deramakot, Tangkulap-Pinangah and Northern Kuamut, n = 1083), in one national park and two nature reserves in Vietnam (Bach Ma National Park, Hue and Quang Nam Saola Nature Reserves, n = 1050), Xe Sap National Park and the ungazetted Ban Palé forest area south of Xe Sap National Park in Laos (n = 165), and Bavarian Forest National Park in Germany (n = 471). A further 30 images of sky with different degrees of cloud cover were taken in Sarawak (Malaysian Borneo) and Germany.
Images were taken with various cameras including Garmin 62 and Garmin Oregon 650 GPS devices and a Panasonic JT-B1 tablet using automatic exposure settings. Images in Bavarian Forest National Park were taken with an AbergBest 21 megapixel digital camera at chest height with the camera leveled using a spirit level on the smartphone. Camera settings were set to automatic mode, no zoom was used.
Masks were created manually in the GNU Image manipulation program (GIMP) using the threshold function. In images of dense tropical canopies with small, overexposed crown gaps

the manual masks tended to show canopy as sky around the crown gaps. These images were selected manually in postprocessing and masks were then created anew in R using value 252 from the red channel for thresholding.

#### Understory vegetation density

Understory vegetation images were taken at the same locations and with the same equipment as the canopy cover images (from Malaysia, Viet Nam and Laos, but not Germany). At each plot, we took four images of a 1x1.5m red flysheet held at a distance of 10m from the camera in each cardinal direction. Images were cropped manually to the extent of the flysheet. Masks were created in GIMP by manually selecting red areas using the Fuzzy Select tool. In total we used 1835 understory vegetation images and their masks in model training and evaluation.

### Data collection recommendations

While overall prediction accuracy was high, overexposed parts of images are uninformative (white) and may thus lead to wrong model predictions. This can happen with bright sunlight shining through canopy gaps, or when photographing directly into the sun. While the models are relatively robust and chances are very high that white regions in input images will be classified as sky, problems can arise if overexposed image regions bleed into neighboring image regions and interfere with their brightness levels (blooming).
To alleviate these problems we recommend taking canopy photos on overcast days, or at least avoid taking images directly into the sun if possible. Ensuring adequate image exposure to avoid overexposure is critical. A slight underexposure of about 1 f-stop is generally recommended for canopy photography. Raw image formats and higher quality cameras can provide higher dynamic range than lower grade cameras. CMOS sensors are less prone to blooming than CCD sensors and can further help avoid problems due to overexposure (Macfarlane *et al.* 2014).

Currently, understory vegetation images are not cropped automatically. To avoid the need for manual cropping we recommend taking pictures in a 8:5 aspect ratio (if unavailable, use 3:2), and zooming in so the red flysheet fills the entire image frame. That method removes the need to crop images and allows direct resizing of images with imageResize(). If this is not possible due to image quality concerns, crop the images using a 5:8 aspect ratio for the cropping mask.
