## Supporting Information S2 for "imageseg: an R package for deep learning-based image segmentation": imageseg_canopy_model_run.R

```
## Canopy density model ##


# R code for creating image segmentation model of forest canopy density
# from R package imageseg

# https://CRAN.R-project.org/package=imageseg 
# https://github.com/EcoDynIZW/imageseg

# see Readme on github for links to model training data


Sys.setenv(RETICULATE_PYTHON = "")

library(imageseg)
library(keras)

# Setup    ####

# directories

# Working directory must contain training data in "~/input/canopy"
dir_main <- getwd()
dir_images  <- file.path(dir_main, "input", "canopy")
dir_model   <- file.path(dir_main, "output", "models")
dirExamples <- file.path(dir_main, "output", "examples", "canopy")

# basic settings

augment <- T   #do data augmentation? TRUE requires a lot of RAM (might exceed 128Gb)

test_split <- 0.2
validation_split <- 0.15

batch_size <- 12
epochs <- 100

optimizer <- "adam"    # "sgd" or "adam"

loss <- "bce_dice"    # "bce_dice" (sum of BCE and soft Dice) or "dice"  (= soft Dice)

dropout <- 0
filters <-  32
layers_per_block <- 2


# callbacks for early stopping and reducing learning rate when validation loss starts decreasing
callbacks <- list(callback_early_stopping(restore_best_weights = TRUE, verbose = 1, patience = 5),    # NULL
                  callback_reduce_lr_on_plateau(factor = 0.2, monitor = "val_loss", patience = 3, verbose = 1))


# Load data ####
lf <- list.files(dir_images, recursive = TRUE, pattern = ".tif$") 
lf_images <- lf[-grep("mask", lf)]
lf_masks <- lf[grep("mask", lf)]


# Test / training split
set.seed(100)
test_index <- sort(sample(1:length(lf_images), length(lf_images) * test_split))


img_test  <- loadImages(fileNames = file.path(dir_images, lf_images[test_index]))
mask_test <- loadImages(fileNames = file.path(dir_images, lf_masks[test_index]))

img_train  <- loadImages(fileNames = file.path(dir_images, lf_images[-test_index]))
mask_train <- loadImages(fileNames = file.path(dir_images, lf_masks[-test_index]))


# Data augmentation
if(augment) {
  img_aug  <- dataAugmentation(images = img_train, rotation_angles = c(0, 90, 180, 270), flip = TRUE, flop = TRUE)
  gc()
  mask_aug <- dataAugmentation(images = mask_train, rotation_angles = c(0, 90, 180, 270), flip = TRUE, flop = TRUE)
  gc()
  
  x_train_all <- imagesToKerasInput(images = img_aug, type = "image")
  gc()
  y_train_all <- imagesToKerasInput(images = mask_aug, type = "mask")
  gc()
} else {
  x_train_all <- imagesToKerasInput(images = img_train, type = "image")
  y_train_all <- imagesToKerasInput(images = mask_train, type = "mask")
}

# convert test images to keras input, without data augmentation
x_test <- imagesToKerasInput(images = img_test, type = "image")
y_test <- imagesToKerasInput(images = mask_test, type = "mask", grayscale = TRUE)
gc()
x_test_info <- img_test$info

# add information about image type to dataframe
x_test_info$type <- NA
x_test_info$type[grep("CCP/", x_test_info$filename)] <- "canopy cover"
x_test_info$type[grep("CCP_sky", x_test_info$filename)] <- "sky"
x_test_info$type[grep("CHP", x_test_info$filename)] <- "canopy closure"


# Validation / training split
validation_index <- sort(sample(1:nrow(x_train_all), round(nrow(x_train_all) * validation_split)))

# subset validation data
x_val <- x_train_all[validation_index,,,]
y_val <- y_train_all[validation_index,,,, drop = FALSE]


# subset training data
x_train <- x_train_all[-validation_index,,,]
y_train <- y_train_all[-validation_index,,,, drop = FALSE]


# validation data created here because we can't define "validation_split" in generator function
# because `validation_split` is only supported for Tensors or NumPy arrays (not generators)
x_train_all_info <- attr(x_train_all, "info")
y_train_all_info <- attr(y_train_all, "info")

# free up memory
rm(x_train_all)
rm(y_train_all)
gc()


# fetch necessary functions
# loss functions
bce_dice_loss  <- imageseg:::bce_dice_loss
dice_coef_loss <- imageseg:::dice_coef_loss      # directly minimize negative Dice coefficient (thus maximize Dice coefficient)

# metrics
dice_coef      <- imageseg:::dice_coef
jaccard_index  <- imageseg:::jaccard_index


# Model training   ####

# create model architecture
model <- u_net(256, 256,
               filters = filters,
               layers_per_block = layers_per_block,
               dropout = dropout
               )


# compile model (with Adam optimizer)
if(optimizer == "adam") {
  if(loss == "bce_dice"){
    model %>% compile(
      optimizer = optimizer_adam(),
      loss      = bce_dice_loss,
      metrics   = list(custom_metric("dice_coef", dice_coef),
                       custom_metric("jaccard_index", jaccard_index)
                      )
    )
  } 
  if(loss == "dice"){
    model %>% compile(
      optimizer = optimizer_adam(),
      loss      = dice_coef_loss,
      metrics   = list(custom_metric("dice_coef", dice_coef),
                       custom_metric("jaccard_index", jaccard_index))
    )
  }
}


# compile model (with SGD optimizer)
if(optimizer == "sgd") {
  learning_rate <- 0.001    # default is 0.01
  decay_rate = learning_rate / epochs
  momentum = 0.8

  model %>% compile(
    optimizer = optimizer_sgd	(lr = learning_rate, decay = decay_rate, momentum = momentum, nesterov = FALSE),
    loss      = lossfun,
    metrics   = list(custom_metric("dice_coef", dice_coef),
                     custom_metric("jaccard_index", jaccard_index))
  )
}


# Generator Function
sampling_generator <- function(x_data, y_data, batch_size) {
  function() {
    rows <- sample(1:nrow(x_data),
                   batch_size,
                   replace = TRUE)
    list(x_data[rows,,,, drop = FALSE], y_data[rows,,,, drop = FALSE])
  }
}


# ca. 9 min/epoch for aug data @256px/ 32filt

history <- model %>% fit(
  x = sampling_generator(x_train, y_train, batch_size = batch_size),
  epochs = epochs,
  steps_per_epoch = round(nrow(x_train) / batch_size),
  validation_data = list(x_val, y_val),
  callbacks = callbacks
)


# model evaluation ####

# overall
scores_all <- model %>% evaluate(x_test, y_test, verbose = 0)

# canopy closure only (hemispherical images)
scores_closure <- model %>% evaluate(x_test[x_test_info$type == "canopy closure",,,, drop = FALSE], 
                                     y_test[x_test_info$type == "canopy closure",,,, drop = FALSE], 
                                     verbose = 0)

# Canopy cover
scores_cover <- model %>% evaluate(x_test[x_test_info$type == "canopy cover",,,, drop = FALSE], 
                                 y_test[x_test_info$type == "canopy cover",,,, drop = FALSE], 
                                 verbose = 0)

# Sky (which is taken as canopy cover images)
scores_sky <- model %>% evaluate(x_test[x_test_info$type == "sky",,,, drop = FALSE], 
                                 y_test[x_test_info$type == "sky",,,, drop = FALSE], 
                                 verbose = 0)
# canopy cover and sky
scores_cover_sky <- model %>% evaluate(x_test[x_test_info$type %in% c("canopy cover", "sky"),,,, drop = FALSE], 
                                     y_test[x_test_info$type %in% c("canopy cover", "sky"),,,, drop = FALSE], 
                                     verbose = 0)


scores_all
scores_closure
scores_cover
scores_sky
scores_cover_sky

# model predictions ####


# only for a few images
n_sample_images <- 200


# separate predictions for CHP and CCP images
x_test_closure   <- x_test[x_test_info$type == "canopy closure",,,, drop = FALSE]
x_test_cover_sky <- x_test[x_test_info$type %in% c("canopy cover", "sky"),,,, drop = FALSE]

prediction_closure <- imageSegmentation(model,
                                        x = x_test_closure[sample(1:nrow(x_test_closure), n_sample_images),,,],
                                        dirExamples = file.path(dirExamples, "closure"),
                                        subsetArea = "circle")        # with circular mask

prediction_cover <- imageSegmentation(model,
                                      x = x_test_cover_sky[sample(1:nrow(x_test_cover_sky), n_sample_images),,,],
                                      dirExamples = file.path(dirExamples, "cover"))

prediction_closure$examples
prediction_cover$examples


# save model
dir.create(dir_model, showWarnings = FALSE)
save_model_hdf5(model, file.path(dir_model, "imageseg_canopy_model.hdf5"))
```
