## Supporting Information S2 for "imageseg: an R package for deep learning-based image segmentation": imageseg_understory_model_run.R

```
## Understory vegetation density model ##


# R code for creating image segmentation model for understory vegetation density
# from R package imageseg

# https://CRAN.R-project.org/package=imageseg 
# https://github.com/EcoDynIZW/imageseg

# see Readme on github for links to model training data


Sys.setenv(RETICULATE_PYTHON = "")

library(imageseg)
library(keras)

# Setup    ####

# directories

# Working directory must contain training data in "~/input/understory"
dir_main <- getwd()
dir_images  <- file.path(dir_main, "input", "understory")
dir_model   <- file.path(dir_main, "output", "models")
dirExamples <- file.path(dir_main, "output", "examples", "understory")


# basic settings for model run
augment <- T   #do data augmentation?

test_split <- 0.2
validation_split <- 0.15

batch_size <- 16
epochs <- 100

optimizer <- "adam"   #  "adam" or  "sgd"

loss <-  "bce_dice"    # "bce_dice" (sum of BCE and soft Dice) or "dice"  (= soft Dice)

dropout <- 0
filters <- 32
layers_per_block <- 2

# callbacks for early stopping and reducing learning rate when validation loss starts decreasing
callbacks <- list(callback_early_stopping(restore_best_weights = TRUE, verbose = 1, patience = ifelse(augment, 5, 10)),    # NULL
                  callback_reduce_lr_on_plateau(factor = 0.2, patience = ifelse(augment, 3, 5), verbose = 1))


# Load data ####
lf <- list.files(dir_images) 
lf_images <- lf[-grep("mask", lf)]
lf_masks <- lf[grep("mask", lf)]


# Test / training split
set.seed(100)
test_index <- sort(sample(1:length(lf_images), length(lf_images) * test_split))


img_test  <- loadImages(fileNames = file.path(dir_images, lf_images[test_index]))
mask_test <- loadImages(fileNames = file.path(dir_images, lf_masks[test_index]))

img_train  <- loadImages(fileNames = file.path(dir_images, lf_images[-test_index]))
mask_train <- loadImages(fileNames = file.path(dir_images, lf_masks[-test_index]))


# Data augmentation
if(augment) {
  img_aug  <- dataAugmentation(images = img_train,  rotation_angles = 0, flop = TRUE)
  mask_aug <- dataAugmentation(images = mask_train, rotation_angles = 0, flop = TRUE)

  x_train_all <- imagesToKerasInput(images = img_aug, type = "image")
  y_train_all <- imagesToKerasInput(images = mask_aug, type = "mask")

} else {
  x_train_all <- imagesToKerasInput(images = img_train, type = "image")
  y_train_all <- imagesToKerasInput(images = mask_train, type = "mask")
}

# convert test images to keras input, without data augmentation
x_test <- imagesToKerasInput(images = img_test, type = "image")
y_test <- imagesToKerasInput(images = mask_test, type = "mask")

x_test_info <- img_test$info

gc()

# Validation / training split
validation_index <- sort(sample(1:nrow(x_train_all), round(nrow(x_train_all) * validation_split)))

# subset validation data
x_val <- x_train_all[validation_index,,,]
y_val <- y_train_all[validation_index,,,, drop = FALSE]

# subset training data
x_train <- x_train_all[-validation_index,,,]
y_train <- y_train_all[-validation_index,,,, drop = FALSE]


# validation data created here because we can't define "validation_split" in generator function
# because `validation_split` is only supported for Tensors or NumPy arrays (not generators)
x_train_all_info <- attr(x_train_all, "info")
y_train_all_info <- attr(y_train_all, "info")

# free up memory
rm(x_train_all)
rm(y_train_all)
gc()


# fetch necessary functions
# loss functions
bce_dice_loss  <- deepForest:::bce_dice_loss       # sum of binary crossentropy and negative Dice coefficient (Soft Dice loss)
dice_coef_loss <- deepForest:::dice_coef_loss      # directly minimize negative Dice coefficient (thus maximize Dice coefficient)

# metrics
dice_coef      <- deepForest:::dice_coef
jaccard_index  <- deepForest:::jaccard_index


# Model training   ####

# create model architecture
model <- u_net(net_w = 160,
               net_h = 256,
               filters = filters,
               layers_per_block = layers_per_block,
               dropout = dropout
               )


# compile model (with Adam optimizer)
if(optimizer == "adam") {
  if(loss == "bce_dice"){
    model %>% compile(
      optimizer = optimizer_adam(),
      loss      = bce_dice_loss,
      metrics   = list(custom_metric("dice_coef", dice_coef),
                       custom_metric("jaccard_index", jaccard_index))
    )
  } 
  if(loss == "dice"){
    model %>% compile(
      optimizer = optimizer_adam(),
      loss      = dice_coef_loss,
      metrics   = list(custom_metric("dice_coef", dice_coef),
                       custom_metric("jaccard_index", jaccard_index))
    )
  }
}


# compile model (with SGD optimizer)
if(optimizer == "sgd") {
  learning_rate <- 0.001    # default is 0.01
  decay_rate = learning_rate / epochs
  momentum = 0.8

  if(loss == "bce"){
    model %>% compile(
      optimizer = optimizer_sgd	(lr = learning_rate, decay = decay_rate, momentum = momentum, nesterov = FALSE),
      loss      =  bce_dice_loss,
      metrics   = list(custom_metric("dice_coef", dice_coef),
                       custom_metric("jaccard_index", jaccard_index))
    )
  } else {
    model %>% compile(
      optimizer = optimizer_sgd	(lr = learning_rate, decay = decay_rate, momentum = momentum, nesterov = FALSE),
      loss      =  dice_coef_loss,
      metrics   = list(custom_metric("dice_coef", dice_coef),
                       custom_metric("jaccard_index", jaccard_index))
    )
  }
}


# Data Generator Function
sampling_generator <- function(x_data, y_data, batch_size) {
  function() {
    rows <- sample(1:nrow(x_data),
                   batch_size,
                   replace = TRUE)
    list(x_data[rows,,,], y_data[rows,,,, drop = FALSE])
  }
}


# fit model
history <- model %>% fit(
  x = sampling_generator(x_train, y_train, batch_size = batch_size),
  epochs = epochs,
  steps_per_epoch = round(nrow(x_train) / batch_size),
  validation_data = list(x_val, y_val),
  callbacks = callbacks
)


# model evaluation ####
scores_all <- model %>% evaluate(x_test, y_test, verbose = 0)

scores_all


# predictions #### 

n_sample_images <- 100

prediction_understory <- imageSegmentation(model,
                            x = x_test[sample(1:nrow(x_test), n_sample_images),,,],
                            dirExamples =  dirExamples)
                            
prediction_understory$examples


# save model
dir.create(dir_model, showWarnings = FALSE)
save_model_hdf5(model, file.path(dir_model, "imageseg_understory_model.hdf5"))
```
